# The general version of Hamilton’s rule

**DOI:** 10.1101/2024.11.25.625193

**Authors:** Matthijs van Veelen

## Abstract

The generality of Hamilton’s rule^1,2^ is much debated^3–14^. In this paper, I show that this debate can be resolved by constructing a general version of Hamilton’s rule, which allows for a large variety of ways in which the fitness of an individual can depend on the social behaviour of oneself and of others. For this, I first derive the Generalized Price equation, which reconnects the Price equation^15^ with the statistics it borrows its terminology from.

The Generalized Price equation moreover shows that there is not just one Price equation, but there is a Price-like equation for every possible true model. This implies that there are also multiple, nested rules to describe selection. The simplest rule is the rule for selection of non-social traits with linear fitness effects. This rule is nested in the classical version of Hamilton’s rule, for which there is consensus that it works for social traits with linear, independent fitness effects. The classical version of Hamilton’s rule, in turn, is nested in more general rules, that for instance allow for nonlinear and/or interdependent fitness effects, like Queller’s rule^16^. The general version of Hamilton’s rule therefore is a constructive solution, that allows us to accurately describe when costly cooperation evolves in a wide variety of circumstances. As a byproduct, we also find a hierarchy of nested rules for selection of non-social traits.

## Introduction

Hamilton’s rule^1,2^ is one of the more famous rules in evolutionary biology. The rule states that altruism will evolve if *rb* > *c*, where *b* is the fitness benefit to the recipient, *c* is the fitness cost to the donor, and *r* is the genetic relatedness between them. The generality of Hamilton’s rule however has always been a topic of contention^3,4^, and positions range all the way from “*Hamilton’s rule almost never holds*”^6^ to “*Inclusive fitness is as general as the genetical theory of natural selection itself*”^7^.

The claim of generality is not based on the original derivation of Hamilton’s rule^1,13,17^, but on later derivations ^8,10,11,18–23^ that use the Price equation^15^. This tradition also started with Hamilton himself^18,19^. The usefulness of the Price equation as a tool for doing theory, however, is not undisputed ^9,12,24–27^. One of the remarkable characteristics of the Price equation literature is that it borrows terms from statistics, like regression coefficients, without also inheriting the natural preoccupations of statistics, such as worrying about model choice or statistical significance.

In this paper, I will show that this debate can be resolved by deriving general versions of the Price equation and of Hamilton’s rule. The Generalized Price equation in regression form generates a set of Price-like equations; one for every different choice for a statistical model, that may describe how the fitness of an individual depends on its genetic makeup. This makes the original Price equation in regression form a special case; it is the Generalized Price equation in regression form, combined with a linear model. The Generalized Price equation repairs the broken link between the Price equation and statistics; when applied to data, standard statistical considerations concerning model choice now translate one-to-one to considerations concerning which of these Price-like equation describes the population genetic dynamics. I will also consider the application of the Generalized Price equation in the modeling domain, and it will be helpful to always discuss the application to data and to modeling separately.

With the Generalized Price equation, I then derive the general version of Hamilton’s rule. Both the limitations of the original Price equation, and ways to overcome these limitations by using the generalized version, are mirrored in the limitations of Hamilton’s rule as we know it, and ways to overcome those. Just like there is not just one Price equation, but a multitude of Price-like equations, there is not just one Hamilton’s rule, but a multitude of Hamilton-like rules. All of them are correct, and all of them are general, but none of them is generally meaningful. A specific Hamilton’s rule, though always correct, is only meaningful if based on the Generalized Price equation, in combination with a model that is appropriately specified for the evolutionary system under study.

The Generalized Price equation puts the debate concerning the validity of Hamilton’s rule in perspective by showing that some arguments are not as decisive as they are currently perceived to be. The side of the debate that claims full generality has repeatedly stressed that Hamilton’s rule, derived with the Price equation, is an identity that holds generally^7,8,10,11^ (that is, it holds, whatever the parent and the offspring generations are).

This is correct, but the general version of Hamilton’s rule shows that being an identity that holds generally cannot be the only relevant characteristic for regarding it as the proper, the right, or the most insightful rule that describes evolution. The reason why this cannot be, is that all Price-like equations generated by the Generalized Price equation, and thereby all Hamilton-like rules produced by them, have those properties; they are all identities that hold with exactly the same generality. To decide between the different Hamilton-like rules, we therefore need additional criteria. When doing empirics, we then have to resort to classical statistics, to see which model agrees with the data. With sufficiently many data, this will then point to a statistical model, and by doing so, it will also point to one of these Hamilton-like rules. An indication that the approach suggested in this paper does indeed pick a Hamilton-like rule that is well-specified, both with modeling and with empirics, is that the quantities in it that we treat as constants are in fact constant, and that they do not change with the composition of the parent population.

## Results

### The Generalized Price equation in combination with models for non-social traits

I would like to start with describing how generalizing the Price equation works. By applying this to linear and non-linear fitness functions in a non-social context, I would moreover like to illustrate that for the Price-like equations that this generates, and that are meant to describe population genetic dynamics, there is scope for under- and for overspecification in the same way that there is scope for under- and for overspecification in statistics.

Assume that we track a set of genes, summarized by a *p*-score^20^. This *p*-score is a value between 0 and 1. In the simplest possible haploid setup, this reflects the presence or absence of a single gene, in which case an individual’s *p*-score would be 1 if the gene is present, and 0 if it is absent. In less simple setups, the *p*-score can for instance reflect a set of genes that affect the same trait. The change in average *p*-score between the parent and the offspring generation can then be written as a sum of two terms, in what Price called the “covariance form” of his equation^15^.

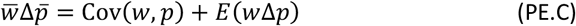

Here, 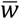 is the average fitness in the parent generation; 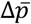 is the change in average *p*-score between parent and offspring generation; 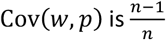 times the sample covariance between fitness and *p*-score (or, in other words, the sample covariance without Bessel’s correction; see Section 2 of Appendix A for details about the terminology); and *E*(*w*Δ*p*) is the fitness-weighted average difference between the *p*-scores of the parents, and, in an asexual model, the *p*-scores of their offspring, or, in a model with sexual reproduction, the *p*-scores of their successful gametes (all details are in Appendix A).

In order to get to the Generalized Price equation in covariance form, we assume a statistical model that has fitness as a dependent variable, and that contains, at least, 1) a constant term, and 2) a linear term for the *p*-score as an explanatory variable. Besides those two terms, the statistical model may contain any number of other terms as well. This statistical model is then combined with the transition we want to apply the Generalized Price equation to. A transition implies that we know what the *p*-score is for all individuals in the parent population, and that we know how many offspring all individuals have, and therefore what their realized fitness is (this is the number of offspring divided by the ploidy). Combining a transition with a statistical model here means that we choose the parameters of the model so that they minimize the sum of squared differences between the model-predicted fitness 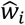 and the realized fitnesses *w*_*i*_, for individuals *i* = 1, …, *n*. For any model that does indeed include a constant term and a linear term for the *p*-score, we can then replace the realized fitnesses *w*_*i*_ in the Price equation with the estimated fitnesses 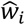 according to the model. This leads to the Generalized Price equation in covariance form:

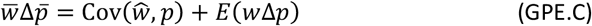

The algebra to show that this is an identity is in Appendix A, on pages 12–14. While replacing *w* with 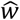 seems like a minimal change, the important thing to keep in mind, is that this equation is an identity for *any* model that contains a constant, and a linear term for the *p*-score. In other words, for any given transition (that is, for any combination of a parent and an offspring population), the model-predicted fitnesses 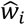 that result from minimizing the sum of squared differences will typically differ, depending on what the model is, but 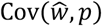 equals the sample covariance *Cov*(*w, p*) for all of these models.

In order to get to what Price calls the “regression form” of the equation, we now choose a set of models for non-social traits, all of which contain a constant and a linear term (which makes it satisfy the condition for the Generalized Price equation to hold). The set of models we choose here is given by

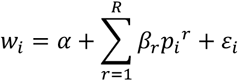

where *w*_*i*_ is the number of offspring of individual *i*; *p*_*i*_ is the *p*-score of individual *i*; *α* is a constant term; *β*_1_,…, *β*_*R*_ are the other coefficients of the model; *ε*_*i*_ is the error term for individual *i*; and where we assume that *R* ≥ 1. This gives a different model for every *R*; a linear model for *R* = 1; a quadratic one for *R* = 2; and so on. If we now minimize the sum of squared differences between the model-predicted fitnesses 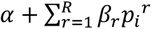 (the right hand side of the equation above, but without the error term) and the realized fitnesses (on the left hand side of the same equation), then we arrive at values 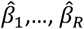 for the regression coefficients.

In Appendix A, on page 15, I show that, for every *R* ≥ 1, and thereby for every model in this set, we can rewrite the Generalized Price equation as follows:

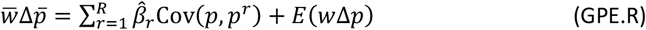

In line with Price’s terminology^15^, this would be the “regression form” of the Generalized Price equation. If we take the linear model (*R* = 1), then, using *Cov(p, p*) = Var(*p*), the above equation becomes

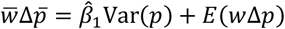

Below we will refer to the linear model as Model A. If instead we take the quadratic model (*R* = 2), the equation becomes

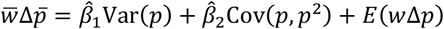

Below we will refer to the quadratic model as Model B. Again, it is important to remember that both of these equations are identities for every possible transition from a parent to an offspring generation. It is also important to recognize that the 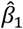 that we arrive at if we combine Model A with a given transition, and the 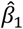 that we arrive at if we use Model B for the same transition, may, and generically will, differ, as we will see.

### The Generalized Price equation applied to two artificial datasets

The following example illustrates how the flexibility of the Generalized Price equation in regression form comes with scope for under- and overspecification. The two Price-like equations above – the one with Model A as a reference model, and the one with Model B as a reference model – are combined with two artificial datasets; one where the offspring generation is generated by Model A; and one where the offspring generation is generated by Model B. Because both equations are identities for all possible transitions between parent and offspring generation, they are also both identities for both of these datasets. Depending on the system under study, the terms in these equations however are not equally meaningful as a description of the population genetic dynamics.

The Generalized Price equation in regression form for Model A, in combination with the offspring generation generated by Model A, returns a value for 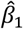 that is very close to the true *β*_1_; 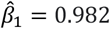, while *β*_1_ = 1. This is depicted in Fig. 1a. All details are in Detailed Calculations A.6 at the end of the Appendices.

**Fig. 1.**
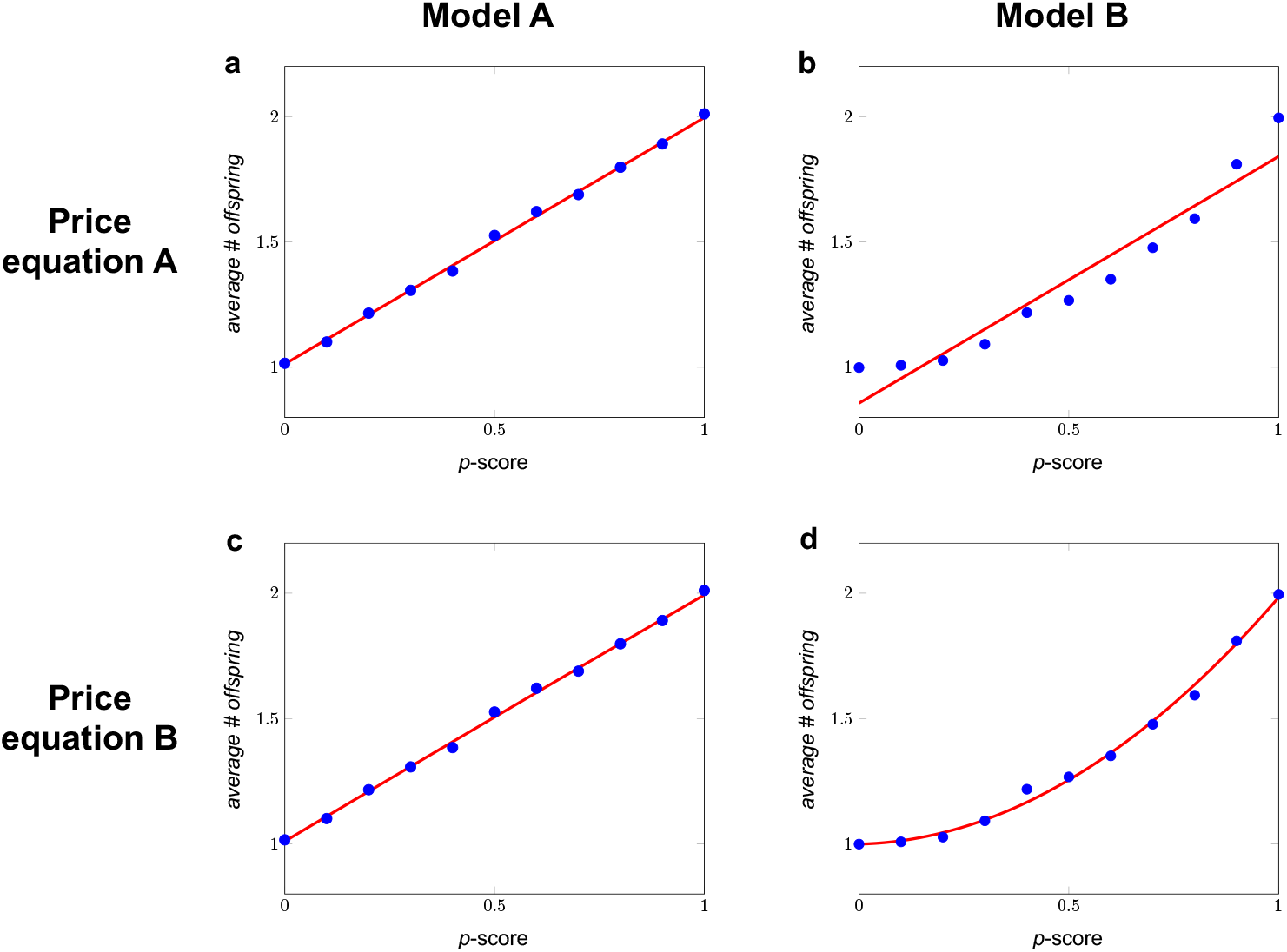
Two Price-like equations and two models. In Model A, the number of offspring follows a binomial distribution with an expected number of offspring of *α* + *β*_1_*p*_*i*_. This means that the model can be summarized as *w*_*i*_ = *α* + *β*_1_*p*_*i*_ + *ε*_*i*_ (see Detailed Calculations A.6 at the end of the Appendices for details). For the transition depicted in panels a and c, we generated an offspring generation using Model A, with *α* = 1 and *β*_1_ = 1. Combining the Generalized Price equation in regression form with Model A includes choosing 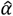 and 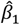 so that they minimize the sum of squared differences between *w*_*i*_ and 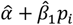. In Model B, the number of offspring follows a binomial distribution with an expected number of offspring of *α* + *β*_1_*p*_*i*_ + *β*_1_*p*_*i*_^2^, which means that *w*_*i*_ = *α* + *β*_1_*p*_*i*_ + *β*_1_*p*_*i*_^2^ + *ε*_*i*_. For the transition depicted in panels b and d, we generated an offspring generation using Model B, with *α* = 1, *β*_1_ = 0, and *β*_2_ = 1. Combining the Generalized Price equation in regression form with Model B includes choosing 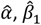, and 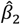 so that they minimize the sum of squared differences between *w*_*i*_ and 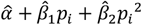. Reproduction is asexual, so parents and their offspring are always identical, and *E*(*w*Δ*p*) = 0 by definition. For both of these models, we started with a population consisting of 2500 parents for each *p*-score, ranging from 0 to 1 in increments of 0.1. The four panels represent the four combinations of the two Price-like equations and the two datasets. The red lines in all panels represent the estimated fitnesses as a function of the *p*-score, as implied by the respective Price-like equations. The aim of this example is not to show that the estimated fitnesses from Price-like equation A match the data generated by Model A, and the estimated fitnesses from Price-like equation B match the data generated by Model B, but the estimated fitnesses from Price equation A do not match the data generated by Model B; that part is obvious. The purpose of this example, instead, is to illustrate that both Price-like equations remain identities, also when they are overspecified (panel c) or underspecified (panel b) with respect to the transition between parent and offspring population they are applied to. With underspecification, it is also visually clear that the estimated fitnesses do *not* match the data, even though Price-like equation A remains an identity, also when combined with data generated by Model B.

The Generalized Price equation in regression form for Model B, in combination with the offspring generation generated by Model B, returns values for 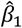 and 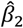 that are also close to their true values; 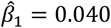, while *β*_1_ = 0; and 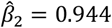, while *β*_1_ = 1. This is depicted in Fig. 1d.

If we combine the Generalized Price equation in regression form for Model B with the offspring generation generated by Model A, then this returns values for 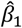 and 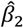 that are also close to their true values; 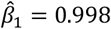, while *β*_1_ = 1; and 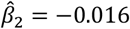, while *β*_2_ = 0. This is depicted in Fig. 1b. Appropriate statistical tests reject that the true *β*_2_ is different from 0, but the harm done by using the Generalized Price equation in regression form for Model B – which is overspecified for data generated by Model A, because it includes a parameter for 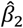, which is really 0 – is therefore limited; the estimated coefficient for *β*_2_ is very close to 0.

The Generalized Price equation in regression form for Model A, in combination with the offspring generation generated by Model B, however, does not return values for 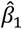 and 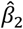 that are close to their true values. For 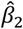 that is obvious, because the statistical model specification sets it to 0, while *β*_1_ = 1. Moreover, 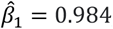, and this is also rather different from the true *β*_1_, which is 0. These differences are the result of underspecification. What is important to note here, is that none of this keeps the Generalized Price equation in regression form for Model A from being an identity, also for data that are really generated by Model B. Being an identity therefore does not guarantee that the regression coefficients in it have a meaningful interpretation; in this case the 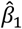 is *not* the linear effect of the *p*-score on fitness.

This also illustrates a general property of the Generalized Price equation. If the data are generated by the model that matches the model that we combine the Generalized Price equation with, then the 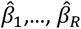 that feature in the regression form are unbiased estimators of the true *β*_1_,…, *β*_*R*_. This is exemplified by panel a, where the Generalized Price equation for Model A is applied to an offspring generation that is really generated by Model A, and by panel d, where the Generalized Price equation for Model B is applied to an offspring generation that is really generated by Model B. If, on the other hand, the model that we combine the Generalized Price equation with is underspecified for the underlying model that generated the data – as it is in the panel b of Fig. 1 – then the regression coefficients are not unbiased estimators of any parameter of the true model. As a symptom of this, if the offspring population is indeed generated by a richer model, then the expected values of the regression coefficients 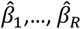 (that is, what values for 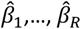 we will find, on average, if we repeatedly draw an offspring population according to the true model) will depend on the composition of the parent population. Again, none of this keeps the Generalized Price equation for Model A from being an identity, also when applied to data generated by Model B.

The regression form of the original Price equation coincides with the regression form of the Generalized Price equation, combined with the linear model for *R* = 1 (Price Equation A in Fig. 1). This implies that limitations of the latter are shared with the original Price equation.

In the Price equation literature, the fact that regression coefficients may not be constant is conflated with, and inadequately labelled as, dynamic insufficiency. We will return to this point after introducing the Generalized Price equation in combination with social traits, but it is worthwhile already here to stress that regression coefficients depending on the composition of the parent population is not a sign of dynamic insufficiency, but rather an indication of misspecification.

### The Generalized Price equation applied in a modeling context

The next example shows that the problem of misspecification is not limited to applications of the Generalized Price equation in a statistical setting. Misspecification is also possible when the Generalized Price equation is applied to a theoretical model. Section 5 in Appendix A describes the conditions under which one can in fact apply the Generalized Price equation in a modeling setting, so here we will assume that those conditions are satisfied; we assume an infinitely large population, error terms with expected value 0, and, if reproduction is sexual, fair meiosis.

In a modeling setting, it seems almost trivial that there is room for misspecification, if we apply the Generalized Price equation for one model to transitions that follow a different model. It will however be useful to acknowledge that this possibility exists. Below, we will therefore consider a linear model, *w*_*i*_ = *α* + *β*_1_*p*_*i*_ + *ε*_*i*_, which we will again refer to as Model A, and a quadratic one, *w*_*i*_ = *α* + *β*_1_*p*_*i*_ + *β*_2_*p*_*i*_^2^ + *ε*_*i*_, which we will refer to as Model B. With infinite populations, such a model then implies that every parent population produces an offspring generation deterministically. If we minimize the squared differences in this transition, with respect to the model that generated the transition, we recover the coefficients of the model exactly (and not with some noise, as in the statistical example above). With two models to combine the Generalized Price equation with, and two models to apply these Price-like equations to, we again have four possible combinations.

As a preparation for moving on to generalizing Hamilton’s rule, I will also point to the rules for selection that the Price-like equations produce.

1. *The Generalized Price equation for Model A applied to transitions following Model A*. The Generalized Price equation in regression form for Model A is

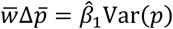 If this equation is applied to a transition that does indeed follow Model A, then the 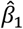 in this Price-like equation is equal to the model parameter *β*_1_. Translating this equation into a criterion for when higher *p*-scores are selected for is straightforward: higher *p*-scores are selected for at all frequencies if and only if 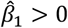. With transitions that are generated by Model A, this means that higher *p*-scores are selected for if and only if *β*_1_ > 0.
2. *The Generalized Price equation for Model B applied to transitions following Model B*. The Generalized Price equation in regression form for Model B is

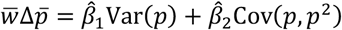 If this equation is applied to a transition that does indeed follow Model B, 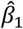 is equal to the model parameter *β*_1_, and 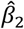 is equal to the model parameter *β*_2_. If we now translate this equation into a criterion for when higher *p*-scores are selected for, then this is still straightforward, but less concise:

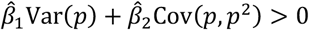 This can be used to describe, for instance, the population genetic dynamics for cases with heterozygote advantage (see Example 3.2 in Appendix A). This is a nice bycatch in the non-social domain of this endeavour, which was meant to solve a long-standing debate in the social domain. If we furthermore assume random mating between generations (see Section 5 of Appendix A), then this simplifies to

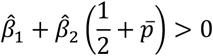

where 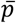 is the average *p*-score in the parent population. With transitions that are indeed generated by Model B, this means that higher *p*-scores are selected for if and only if 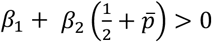. The rule for the Generalized Price equation in regression form for Model A, applied to transitions that follow Model A, is nested in this rule here, in the sense that if we choose *β*_2_ = 0, this rule reverts to *β*_1_ > 0.
3. *The Generalized Price equation for Model B applied to transitions following Model A* The Generalized Price equation in regression form for Model B is still

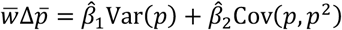 If this equation is applied to a transition that follows Model A, then 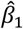 is equal to the model parameter *β*_1_, and 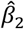 is equal to the model parameter *β*_2_, which is 0. The overspecification therefore is inconsequential; we end up with the same equation that we arrived at when we applied the Generalized Price equation in regression form for Model A to a transition that follows Model A.
4. *The Generalized Price equation for Model A applied to transitions following Model B*

The Generalized Price equation in regression form for Model A is still

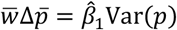

If applied to a transition that follows Model B, then this implies that 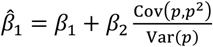.

What is important to realize, is that now 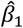 is no longer a constant, as it depends on the population state through the 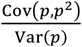-term. Also, it would be wrong to interpret 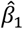 as the linear effect of the gene on fitness, because here 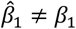.

If we translate this equation into a criterion for when higher *p*-scores are is selected for, then this would be the same rule as the one we arrived at when the Generalized Price equation in regression form for Model A was applied to Model A, which is that higher *p*-scores are selected for if 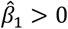. This is still a rule, and it gets the direction of selection right, but the 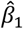 in it does not have a meaningful interpretation anymore. The fact that the regression coefficient 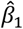 here changes with the population state moreover reflects misspecification, but now in a modeling context, and also here this does not reflect dynamic insufficiency.

### The general version of Hamilton’s rule

For the derivation of the general version of Hamilton’s rule, we need a richer setup, with two variables instead of one: a *p*-score and a *q*-score. The *p*-score represents a set of genes that code for some social behaviour, and the *q*-score of an individual represents the *p*-score of its interaction partner. We assume that both can take values in [0,1]. We will combine the Generalized Price equation in regression form with a general set of models for social traits. This requires some formal notation.

Here we choose a set of models, in which a model is defined by its non-zero coefficients. That is, a model is a choice for a set of non-zero coefficients *E*, where (*k, l*) ∈ *E* if the term *p*_*i*_^*k*^*q*_*i*_^*l*^ in the model has a non-zero coefficient *β*_*k,l*_. This then defines a model as follows:

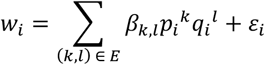

where *ε*_*i*_ is a noise term with expected value 0. We assume that coefficients *β*_0,0_ and *β*_1,0_ are included, or, in other words, that the model contains a constant and a linear term for the *p*-score.

We can take any such model as a reference model for the Generalized Price equation in regression form. If we do, we find the following Price-like equation for the population genetic dynamics (see Section 4 of Appendix A for details).

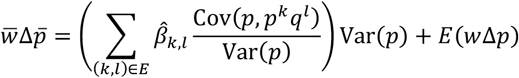

If we define *E*′ as the set of non-zero coefficients of the model, besides the constant term and the linear term for the *p*-score, then the generalized version of Hamilton’s rule is that 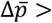 if and only if

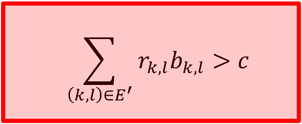

where *c* = −*β*_1,0_, *bk,l* = *β*_*k,l*_, and 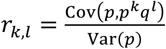, both for all (*k, l*) ∈ *E*′. The linear term for the *p*-score still features in this equation, because that is what *c* represents. Details are in Section 3 of Appendix B. In Section 5 of Appendix B this is generalized further to include interactions between more than two individuals.

The general version of Hamilton’s rule produces a set of rules; one for every model of how traits affect fitnesses. Less general models, that are nested in more general ones, come with rules that are nested in more general rules. This includes rules for non-social traits. We get the rules for non-social traits, if the set of model coefficients, besides a constant, only includes coefficients *β*_*k*,0_, and no coefficients for terms that relate to the *q*-score. The simplest such rule is the linear one that we have seen above, and that we get if the true model only contains a constant, and a term for the linear effect of the *p*-score. Above, we referred to this model as Model A. This rule is nested in the rule for the quadratic model, which has a constant, a term for the linear effect of the *p*-score, and a term for the quadratic effect of the *p*-score. We referred to this model as Model B above. This describes selection in case of, for instance, heterozygote advantage (see Appendix A, Sections 3 and 5). This rule, in turn, is nested in more general rules for selection of non-social traits that include coefficients *β*_*k*,0_ for *k* > 2.

If we then go back to the rule for non-social traits, but with linear fitness effects only, then this rule is also nested in the classical Hamilton’s rule. The classical Hamilton’s rule is the rule that we get for a model that includes a constant, a term for the linear effect of the *p*-score, and a term for the linear effect of the *q*-score. This rule, in turn, is nested in the rule that we get when an interaction effect between the *p*- and the *q*-score is added to the model. The classical Hamilton’s rule therefore is itself nested in what is best refered to as Queller’s rule^16^. These rules moreover allow for further nesting, if the model we are interested in is more general, or if the data suggest that the true model includes additional non-zero coefficients. Below, we will however restrict attention to these three nested models in order to illustrate how they produce three rules that are nested, and how using the generalized version of Hamilton’s rule allows us to see the debate on the generality of the classical Hamilton’s rule in a new light.

### Three models and three rules

In Model 1, the *p*-score relates to a non-social trait, that only affects the fitness of its carrier. We also referred to this model as Model A before, where we nested it in a different, also non-social, but more general model.

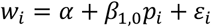

In Model 2, the *p*-score relates to a social trait, that affects the fitness of its carrier, and the fitness of the partner of its carrier. In the fitness function, this is reflected by the effect of the *p*-score of the partner (the *q*-score) on individual *i*. These effects are moreover assumed to be independent; the effect on the partner is the same, irrespective of the *p*-score of the partner.

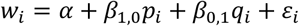

In Model 3, the *p*-score also relates to a social trait, but now the effects are not assumed to be independent; the effect on the partner may depend on the *p*-score of the partner. This is reflected by the interaction term.

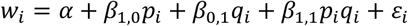

#### Model 1: selection of a non-social trait

The Generalized Price equation in regression form for Model 1 is

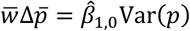

The rule that this Price-like equation produces is that higher *p*-scores will be selected for if and only if 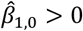. This is also the rule that we found above, when we referred to Model 1 as Model A. If the model under consideration is indeed Model 1, then 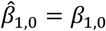, both in the Price-like equation, and in the rule for selection.

#### Model 2: selection of a linear social trait

The Generalized Price equation in regression form for Model 2 is

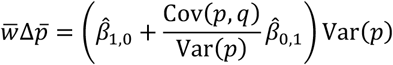

From this, we can see that 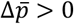 if and only if 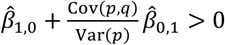. If the model under consideration is indeed Model 2, then, with an infinitely large population, the regression coefficients 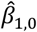 and 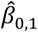 will coincide with the true values of the parameters *β*_1,0_ and *β*_0,1_, respectively, and 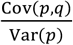 will coincide with relatedness *r* between the individuals that interact. If we then define *c* = −*β*_1,0_ and *b* = *β*_0,1_, we can rewrite this as

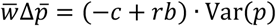

Here we naturally recognize the classical Hamilton’s rule, because this implies that 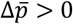 if and only if *rb* > *c*.

At this point, it is useful to observe that if this is a social trait, and the true model is Model 2, but we combine it with the Generalized Price equation in regression form for Model 1, the latter still gets the direction of selection right. In that case, we would still use

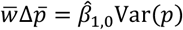

but now with 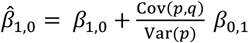. This however is typically not how we describe the evolution of social traits. The reason why we do not describe selection of a social trait with the rule for the non-social model is however *not* that this rule would get the direction of selection wrong; it would not. As a matter of fact, this rule always gets the direction of selection right too. The reason that here we use the classical version of Hamilton’s rule instead, is that the classical version of Hamilton’s rule reveals the actual population genetic dynamics that govern selection; with a social trait for which *rb* > *c*, the reason why the genes for the trait are selected is not that the behaviour they induce is good for the fitness of the individual that carries these genes itself; the reason is that the fitness costs of the associated behaviour to the individual itself are outweighed by how much fitness benefits carriers get through related individuals, compared to how much such benefits non-carriers get. In a theory model, this is postulated. With data, and if the sample size is large enough, statistical tests would make the same determination, by pointing to Model 2, and not Model 1, as the true model, if it is in fact the true model.

At this point, we therefore have two Price-like equations: the one for Model 1 and the one for Model 2. Both of these Price-like equations are identities, and both of them are general. When applied to Model 2, however, the Price-like equation for Model 1 is not meaningful, in the sense that the regression coefficient 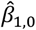 does not reflect the effect of the behaviour on the individual itself. As a symptom of this, it is not a constant, and varies with the population state. The core point of the original derivation of Hamilton’s rule, using the original Price equation, therefore is to not use an underspecified model. It is useful to keep this in mind when we are faced with an analogous choice between Price-like equations, and between rules based on them, below.

#### Model 3: selection of a social trait with an interaction effect

The Generalized Price equation in regression form for the third model is

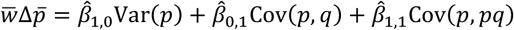

If the model under consideration is indeed Model 3, and we assume an infinite population, the regression coefficients 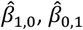, and 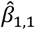 will coincide with the true values of the parameters *β*_1,0_, *β*_0,1_, and *β*_1,1_, respectively. Also, in an infinite population model, 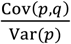 will coincide with relatedness *r*, which we call *r*_0,1_ here, in order to distinguish it from 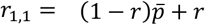. The latter is what 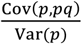 will be in an infinite population (see Appendix B). If we then define *c* = −*β*_1,0_, *b*_0,1_ = *β*_0,1_, and *b*_1,1_ = *β*_1,1_, we can rewrite the Generalized Price equation in regression form for the third model as

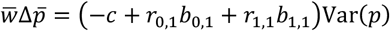

This does not give us the classical Hamilton’s rule that we are familiar with. It does however give us a correct criterion for when higher *p*-scores are selected for; 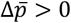 if and only if *r*_0,1_*b*_0,1_ + *r*_1,1_*b*_1,1_ > *c*. This criterion features two benefits and two relatednesses; besides the classical, linear benefit *b*, which we call *b*_0,1_ here, it also has an interaction benefit *b*_1,1_, that only fully materializes if both the individual itself and its partner have a *p*-score of 1. If *b*_1,1_ = 0, then we are back in Model 2, and this rule reverts to the classical Hamilton’s rule. The rule with a non-zero interaction effect is the same as the rule suggested by Queller^16^, if we assume that genotype and phenotype correlate perfectly (see also Section 2 of Appendix B).

We can, of course, still apply the Generalized Price equation in regression form for Model 1, or the Generalized Price equation for Model 2, to Model 3 in the same way that we can apply the Generalized Price equation for Model 1 to Model 2, as we did above. We do this in detail in Section 2 of Appendix B to make sure that there can be no misunderstanding concerning the details, and the results are shown in Figure 2. The important observation to make here is that both of these rules *also* get the direction of selection right. The argument for not using these rules for Model 3, even though they do get the direction of selection right, is identical to the argument for not using the rule for selection of non-social traits for social traits: the rule for non-social traits just does not describe the population genetic dynamics for a social trait accurately. This is why we use the conventional version of Hamilton’s rule when modelling social behaviour, and not the rule that we would get if we applied the Generalized Price equation in regression form for the non-social model (Model 1). The Price-like equation for the non-social model also gets the direction of selection right, but we know that this version is incomplete, precisely because it leaves out the true social effects we care about unveiling when we use the conventional Hamilton’s rule. For exactly analogous reasons, we should not rely on underspecified versions of the Price equation in other instances, including the case where the conventional Hamilton’s rule itself, in turn, is based on an underspecified Price-like equation.

**Fig. 2.**
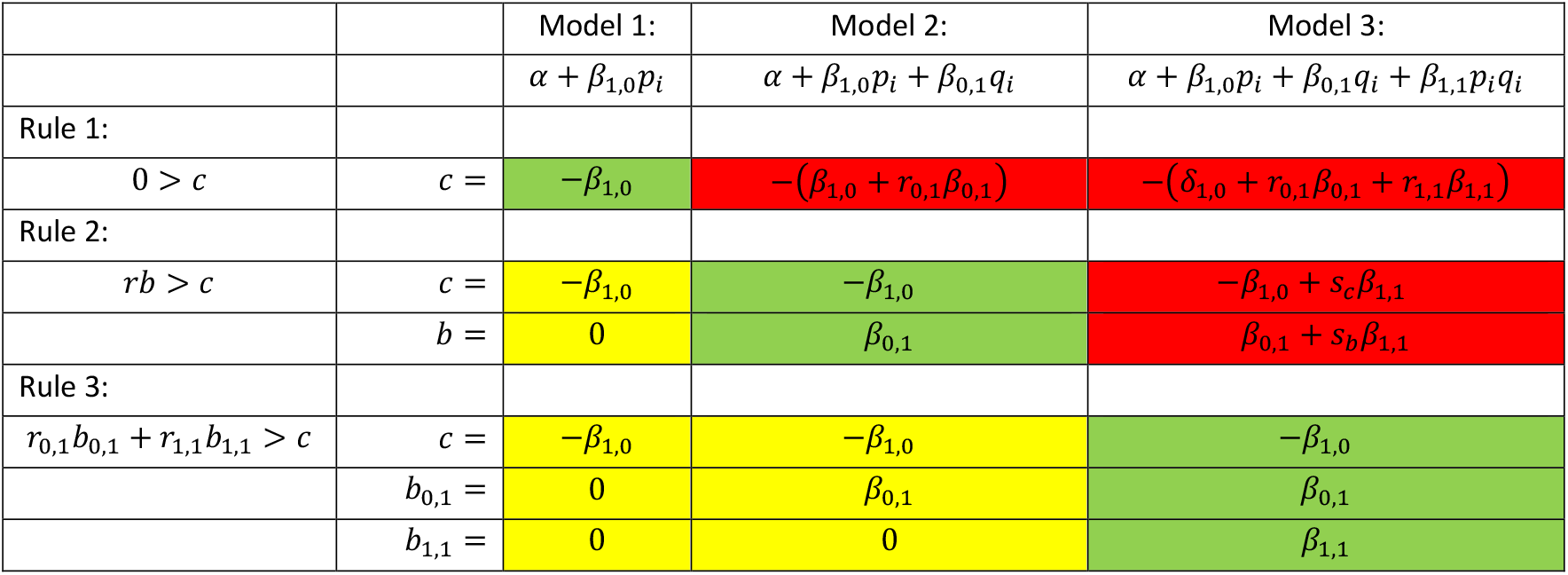
Three rules and three models. This table gives all combinations of the three rules and the three models discussed in the example. All rules indicate the direction of selection correctly for all models. Yellow indicates a combination of a rule and a model, where the rule is more general than is needed for the model. This leads to one or more *b*’s being 0. These are relatively harmless overspecifications. Red indicates a combination of a rule and a model, where the rule is not general enough (underspecified) for the model. This leads to *b*’s and *c*’s that depend on the population state. Terms that depend on the population state are abbreviated as follows: 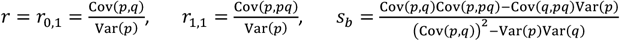 and 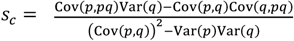 (see Detailed Calculations B.3 at the end of the Appendices for calculations). Rule 1 is the standard rule for non-social evolution for linear non-social traits. Rule 2 is the classical Hamilton’s rule. Rule 3 is Queller’s rule^16^, which is a rule that allows for an interaction effect. Rule 1 is nested in Rule 2, which is nested in Rule 3, which can be nested in a more general rules as well.

As a symptom of underspecification, if we apply the Generalized Price equation in regression form for Model 1 to Model 3, then the value for 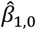 is not a constant, and depends on the population state. Similarly, if we apply the Generalized Price equation for Model 2 to Model 3, then the values for 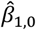 and 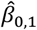 are not constants, and depend on the population state.

The part of the literature on Hamilton’s rule that has emphasized its full generality, has however insisted that we always use one and the same equation, even if it is underspecified, and incorporate more general models, such as Model 3, within the classical version of Hamilton’s rule^7,8,10,11,20,23,28,29^.

The relation between these examples can be summarized in a table that, for the three models and the three rules, represents all nine combinations of them (see Fig. 2). This matrix of combinations of models and rules also helps to understand what causes the contentiousness of the debate on the generality of Hamilton’s rule. Rule 2, which is the rule that we get from applying the standard Price equation (also known as using the regression method), is a completely general rule, in the sense that whatever the true model is, it always gets the direction of selection right. This is true, but that fact is not a good argument for singling this rule out as more helpful, meaningful, or insightful than other rules. In the example above, we have seen that if we apply the Generalized Price equation in regression form (which one could also describe as using the regression method, but now with a richer menu of alternative underlying statistical models) then this can also give us Rule 1 or Rule 3, depending on the statistical reference model we use. These rules are equally correct, in the sense that they also always get the direction of selection right. They are also equally general, in the sense that they also get it right for every possible model. Being a general rule, that always gets the direction of selection right, therefore, cannot be a criterion for elevating Rule 2 (which is the classical Hamilton’s rule), above the other ones, because Rules 1 and 3 are also general rules, that always get the direction of selection right.

Since being completely general and always getting the direction of selection right does not single out any of the possible rules, we need additional criteria. A natural criterion would be that besides being correct, the terms in the rule would have to be meaningful. More precisely, we think the rule should do what Rule 1 (the rule for evolution of non-social traits) does for Model 1, and what Rule 2 (the classical Hamilton’s rule) does for Model 2, and that is to separate model parameters from properties of the population state, in order to produce an equation that describes the population genetic dynamics. That is what Rule 3 does for Model 3. Ever more general models would moreover require ever more general rules to accurately capture the population dynamics.

The debate in the literature on the generality of Hamilton’s rule is so long-lasting because it mostly focuses on whether or not rules are correct, and not on whether the rules are (also) meaningful. One side of the debate tends to return to the argument that Rule 2 is general and correct^7,8,10,11^. The other side of the debate tends to return to models that do not fit Model 2^3–5^. Sometimes arguments on this side take a completely different approach, and rather than describing selection with the Price equation, and then worry about whether or not one can interpret regression coefficients in it as benefits and costs, they start with models with a priori interpretable definitions for *b* and *c*. This is called the counterfactual approach to defining costs and benefits^13,14^, as opposed to the regression approach, and because it has a different definition of the costs and benefits, it can result in rules that end up getting the direction of selection wrong^3,4,13,14^.

Another recurrent point is that when regression coefficients in the Price equation are found to vary with the state of the parent population, this is often claimed to be the result of the Price equation being dynamically insufficient. In order to point out why that is inaccurate, we can first point to ways in which an underlying model *can* be dynamically insufficient. If we have a model that describes how the fitness of an individual depends on its *p*- and *q*-score, and we are given a parent population, then this model would produce the *p*-scores in the new generation. It would however not generate which individuals are partnered up with whom in the new generation, and therefore it would not identify who has which *q*-score in the new generation. In this case a full model would have to include more than just the fitness function, but in the absense of assumptions about the matching in the new generation, such a model would be dynamically insufficient. It is however important to notice that this is *not* the same as the regression coefficients 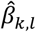, depending on the population state, which is regularly also referred to as dynamical insufficiency. Describing this as dynamical insufficiency is incorrect, and the dependence on the population state is really a symptom of misspecification. Section 6 of Appendix A elaborates on this point in a more precise way^30^.

## Discussion

The general version of Hamilton’s rule provides a positive resolution of a long-standing, heated debate. I derived the Generalized Price equation, which reconnects the Price equation with statistics, and allows us to base rules for selection properly on dynamical models for population genetics that offer the flexibility needed to accommodate a variety of ways in which fitness can depend on traits of individuals and their interaction partners.

While previous work focused on the limitations of the original Price equation^24,25,27^, and on limitations of the classical version of Hamilton’s rule, when there are interaction effects ^3,4,6,9,12–14^, the Generalized Price equation, and the general version of Hamilton’s rule that is derived with it, are constructive contributions that offer alternatives for a large variety of forms the fitness function can have. The insight that comes with it has major implications, also for empirical research on kin selection. It implies that whether or not Hamilton’s rule holds, is not a meaningful empirical question. As a field, we have nonetheless spent quite some time and energy on trying to answer this question^31^. Because the question turns out to be ill-posed, we can stop trying to answer it. The meaningful empirical question that we should focus on instead, is what form the fitness function has, and which version of Hamilton’s rule therefore applies to which social trait.

## Notes

### Competing Interest Statement

The authors have declared no competing interest.

### Summary of Updates

I made a few small edits to improve the text.

